# *in vivo*-like Scaffold-free 3D *in vitro* Models of Muscular Dystrophies: The Case for Anchored Cell Sheet Engineering in Personalized Medicine

**DOI:** 10.1101/2024.11.16.623949

**Authors:** Alireza Shahin-Shamsabadi, John Cappuccitti

## Abstract

Progress in understanding the underlying mechanisms of muscle dystrophies and finding effective treatments for them has been hindered by the absence of relevant *in vitro* models for biomedical research. In this study, an entirely scaffold-free cell sheet engineering-based platform is used to make such *in vitro* models using patient-specific cells. Unlike reductionist bottom-up approaches, this holistic biofabrication method, termed anchored cell sheet engineering, effectively replicated mature cell phenotypes and tissue- and disease-specific ECM deposited by the cells themselves. Robust anchored 3D muscle fibers were developed using primary cells from both healthy individuals and patients with Duchenne dystrophy and Myotonic dystrophy type 1. Through a combination of histology, immunostaining, and proteomics analysis, it was demonstrated that these models formed mature constructs that closely resembled *in vivo* conditions, outperforming traditional 2D cultures in their translation potential. Models of diseased tissues, analyzed through various analysis, accurately reflected key phenotypic features of the respective diseases. Furthermore, when treated with therapeutically beneficial drugs, the detailed changes in their proteomic profiles were documented. This novel *in vitro* modeling approach, compared to other 3D techniques that use exogenous scaffolding or bioink, provides a promising platform for advancing the development of muscle dystrophy models, among other conditions.

## 1. Introduction

Muscular dystrophies, a heterogeneous group of inherited myogenic disorders, profoundly impact patient quality of life through progressive muscle degradation and weakness, with current therapeutic interventions offering limited but promising approaches to modify disease progression in specific dystrophy subtypes [1, 2]. Myotonic dystrophy and Duchenne muscular dystrophy are the two most prevalent dystrophies in adults. Myotonic dystrophy, an autosomal dominant disorder, manifests with multisystemic symptoms such as muscle weakness, myotonia, and cardiac abnormalities [3] and Duchenne muscular dystrophy is characterized by progressive muscle degeneration and weakness [4]. In these muscular dystrophies, muscle weakness is the primary symptom, but clinical and pathologic criteria, as well as genetic basis, have been used for diagnostic and classification purposes. The knowledge regarding muscular dystrophies is expanding, however, even with significant research, therapeutic options remain limited and largely palliative. Currently available treatments for both of these dystrophies focus primarily on managing symptoms rather than addressing the underlying disease mechanisms. In this context, robust models that can more accurately mimic individual patients’ disease pathology are crucial for the development of novel therapeutics [5, 6].

Animal models, such as rodent, porcine, and canine, have provided valuable insights into the mechanisms underlying dystrophies and potential approaches for therapeutic interventions. However, many of the genes involved in these dystrophies do not exist in any of the available animal models [7]. Furthermore, the use of animal studies has been constrained by both ethical considerations and high costs [8]. Many available diagnostic and therapeutic strategies have proven to be mutation-specific, underscoring the increasing need for a personalized approach towards the detection and treatment of these diseases [2, 4, 9]. To address the limitations of animal models for muscular dystrophies and to fulfill the demands of personalized treatments, development of *in vitro* models that closely replicate *in vivo* tissue characteristics has become increasingly important [6]. Regardless of the origin and cause of these dystrophies, the cells that form the tissue are one of the major components responsible for the development and progression of the dystrophy. While various muscle-resident cell populations, including fibroblasts, pericytes, endothelial cells, neural cells, and immune system cells, play critical roles in these diseases, tissue-specific skeletal muscle cells are central to maintaining tissue homeostasis and driving disease progression. The cell source used to develop relevant *in vitro* models should ideally exhibit proper proliferation and the ability to differentiate and form mature tissues [10]. A variety of *in vitro* models for these diseases have been developed using patient-derived induced pluripotent stem cells (iPSCs) [11], primary progenitor cell cultures [12], and even immortalized muscle cells [13]. Each cell type presents its own set of advantages and limitations, with iPSC models offering genetic representativeness and primary cultures providing better cell maturity.

In addition to the choice of cell type and source for making relevant *in vitro* models, the culture system also plays a major role in recreating *in vivo*-like microenvironments. While traditional two-dimensional (2D) cell culture systems have long been foundational in cellular studies because of their reproducibility and ease of use, their oversimplified nature limits their ability to replicate the complex cellular microenvironments and interactions found in living organisms. This limitation often results in discrepancies between *in vitro* findings and clinical outcomes. underscoring the necessity for more physiologically relevant three-dimensional (3D) models, such as organoids and organ-on-chip systems, that can offer more accurate platforms for drug discovery and disease modeling [14, 15].

To accurately reflect the genetic background, origin, and mechanisms of various muscular dystrophies, and to authentically simulate their development and progression, *in vitro* models need to properly incorporate both the cells and their microenvironment, predominantly defined by the tissue’s extracellular matrix (ECM) [10]. The role of ECM disruption in many of these dystrophies and the need for recreating such aspects of the disease for personalized *in vitro* models has been shown before [16]. Many of these studies employ tissue engineering techniques using various biomaterials, however, these exogenous and unfamiliar materials lack the appropriate biochemical composition and mechanical cues that affect cells’ morphology, adhesion, proliferation, and phenotype maintenance, thereby compromising the accuracy of the models [10]. Attempts have also been made to use ECM-derived hydrogels to more closely mimic the *in vivo* characteristics of skeletal muscle tissue [17]. However, even these biomaterials fail to fully meet the specific requirements necessary to recreate disease conditions accurately.

As an alternative to techniques that rely on exogenous biomaterials for scaffolds or bioinks, scaffold-free 3D culture techniques induce cells to secrete and preserve their own ECM. These approaches enhance cell- cell and cell-matrix interactions, which are crucial for replicating physiologically relevant microenvironments [18, 19]. Among these scaffold-free techniques, cell sheet engineering has shown promise [20, 21] and has been used for recreating *in vivo*-like skeletal muscle constructs as well. However, these constructs have been limited to sheet-like formats, and without the external anchors necessary to maintain their integrity, their application in *in vitro* modeling has not been widespread [22, 23].

To overcome these limitations, a recent innovation introduced a cell sheet engineering-based platform capable of biofabricating 3D, scaffold-free, and anchored constructs in any formfactor. This platform has proven capable of recreating *in vitro* tissues that exhibit superior differentiation and mature cell phenotypes, along with accurate ECM content. Here, this novel platform, called anchored cell sheet engineering [24], is used to develop patient-specific *in vitro* models of skeletal muscle. These models used primary cells sourced from both healthy individuals and individuals with Duchenne and Myotonic dystrophies. By using a combination of histological and immunostaining analysis alongside comprehensive proteomics, it was demonstrated that these anchored cell sheet engineering-based 3D muscle fibers significantly outperformed traditional 2D cultures in replicating the structural and biochemical properties of *in vivo* muscle tissues. It was shown that anchored cell sheet engineering-based 3D models of Duchenne and Myotonic dystrophy, compared to healthy counterparts, exhibit protein expressions and both upregulated and downregulated pathways that closely mirror *in vivo* observations. By administering drug treatments to these diseased models, the exact mechanisms of action for modulating protein expression patterns were understood, providing a clear path toward more effective therapeutic strategies. This work not only underscores the inadequacies of conventional 2D models, but also illustrates the potential of anchored cell sheet engineering as a transformative tool for the mechanistic study of muscular dystrophies and the acceleration of drug discovery processes. As this biofabrication platform undergoes further refinement, a more robust platform capable of contributing to the development of curative therapies will be generated. This advancement could significantly transform the treatment landscape for debilitating muscle diseases and other conditions.

## 2. Results and Discussion

### Biofabrication of 3D anchored muscle fibers for disease modeling

Anchored cell sheet engineering [24], a scaffold-free biofabrication platform based on the cell sheet engineering concept, was used to develop 3D *in vitro* models of skeletal muscle tissue. Briefly, culture devices were fabricated using a 3D printed master mold with the inverse design of the culture device. Different resins, including Ecoflex 00-30, polydimethylsiloxane (PDMS), and a mixture of PDMS and Mold Star™ 15, were casted as pillars, base of the culture device, and culture device walls, respectively. The parallel patterns of 3D printing surface texture, consisting of shallow grooves, were replicated on the PDMS base membrane and were used to align the cells and induce them to deposit higher ECM content. The hydrophilicity of the PDMS base membrane was adjusted through treatment with tannic acid solution followed by autoclaving for sterilization. Cell culture was started immediately after treating the culture devices with vitronectin to improve cell attachment.

For cell sheet engineering purposes, each culture device was secured on a sterile 3D printed support (**Figure 1a**). On day one, culture was started by adding 5×10^5^ primary myoblasts in 3 mL of growth medium. On day 3, with cells at near 100% confluency, the medium was switched to their differentiation medium with 100 µg/mL 2-Phospho-L-ascorbic acid trisodium. This medium was refreshed once every three days followed by the addition of 5×10^5^ cells every 3 days starting on day 4 until day 19. On day 20 a multi-layered cell and ECM construct had formed. This layer, now loosely attached to the membrane, was released from the membrane by gently scraping it from four edges of the culture device using a cell scraper. After 24 hours, the cell sheet released itself completely from the membrane and remodeled itself to form a 3D anchored fiber, recognizing the pillars as new anchors. After an additional 72 hours in culture, fibers had become completely mature (**Figures 1b and c**). The overall process is shown in **Figure 1d**. It was shown in the previous study that 3D printing surface textures created parallel patterns on the PDMS surface at different scales, including mico- and meso-scales, and were sufficient for skeletal muscle cells to be fully aligned. It was also shown that the presence of these alignments as well as shallow grooves increased cells’ ECM production and their maturation levels [24].

**Figure 1.**
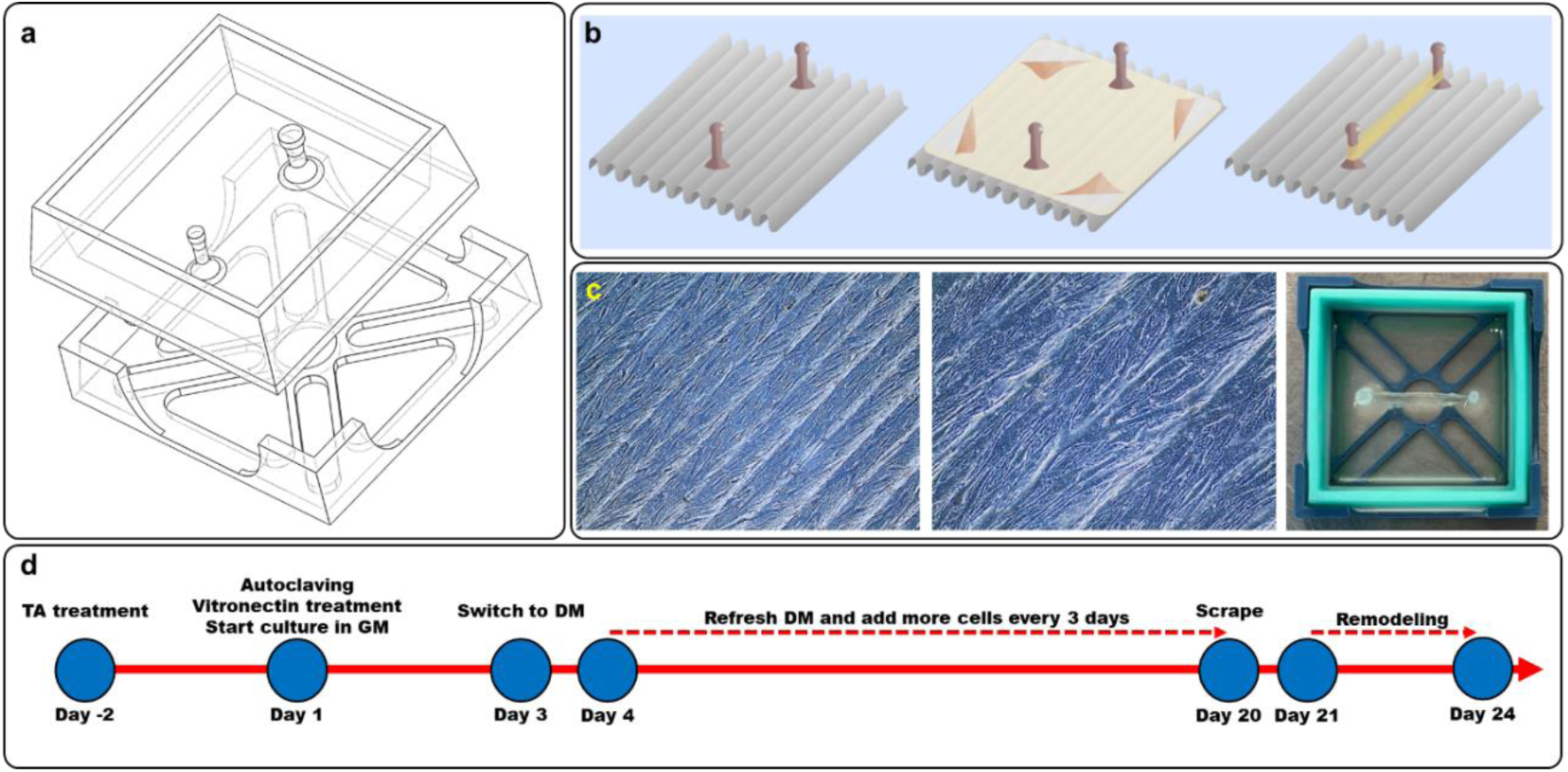
**a)** Schematic of culture device and its support, **b)** schematic of anchored cell sheet engineering process, **c)** alignment and differentiation of skeletal muscle cells on patterned membrane and eventual 3D anchored fiber formation, **d)** overall process showing culture device treatment followed by 2D and 3D cell cultures. 3D printing surface patterns are replicated on the PDMS membrane aligning the cells and inducing them to deposit relevant ECM content. After growing multiple layers of cells on the membrane, the cell layer’s attachment to the PDMS membrane weakens and scraping the edges of the layer from the membrane facilitates its complete lift-off. The detached cell sheet understands the presence of pillars and uses them as anchors to guide its remodeling into a mature, anchored 3D muscle fiber. TA: tannic acid, GM: growth medium, DM: differentiation medium.

### Evaluation of cellular differentiation and maturation in scaffold-free 3D anchored fibers

The 3D anchored fibers were biofabricated using cells from a healthy subject (here called HC group) and two others, one with Myotonic dystrophy type 1 (here called MD group) and another with Duchenne muscular dystrophy (here called DD group). Commercially available cells from CookMyosite® were used for this purpose (**Table 1**). Cells were grown in their growth medium and, after replating them to the culture devices and reaching 100% confluency, were differentiated using their differentiation medium supplemented with 100 µg/mL 2-Phospho-L-ascorbic acid trisodium for improved ECM production. Formation of differentiated and mature 3D scaffold-free anchored fibers using all three cell types was confirmed through immunostaining by targeting for desmin and myosin heavy chain (MyHC) (**Figure 2**). MyHC is a major contractile protein of skeletal muscle tissue, appearing in early stages of differentiation as myoblasts fuse to form myotubes. Desmin, an intermediate filament protein, emerges during the alignment and fusion of myoblasts, playing a crucial role in organizing the cytoskeleton and maintaining the muscle structural stability. Its expression increases as the muscle fibers mature [25–27].

**Figure 2.**
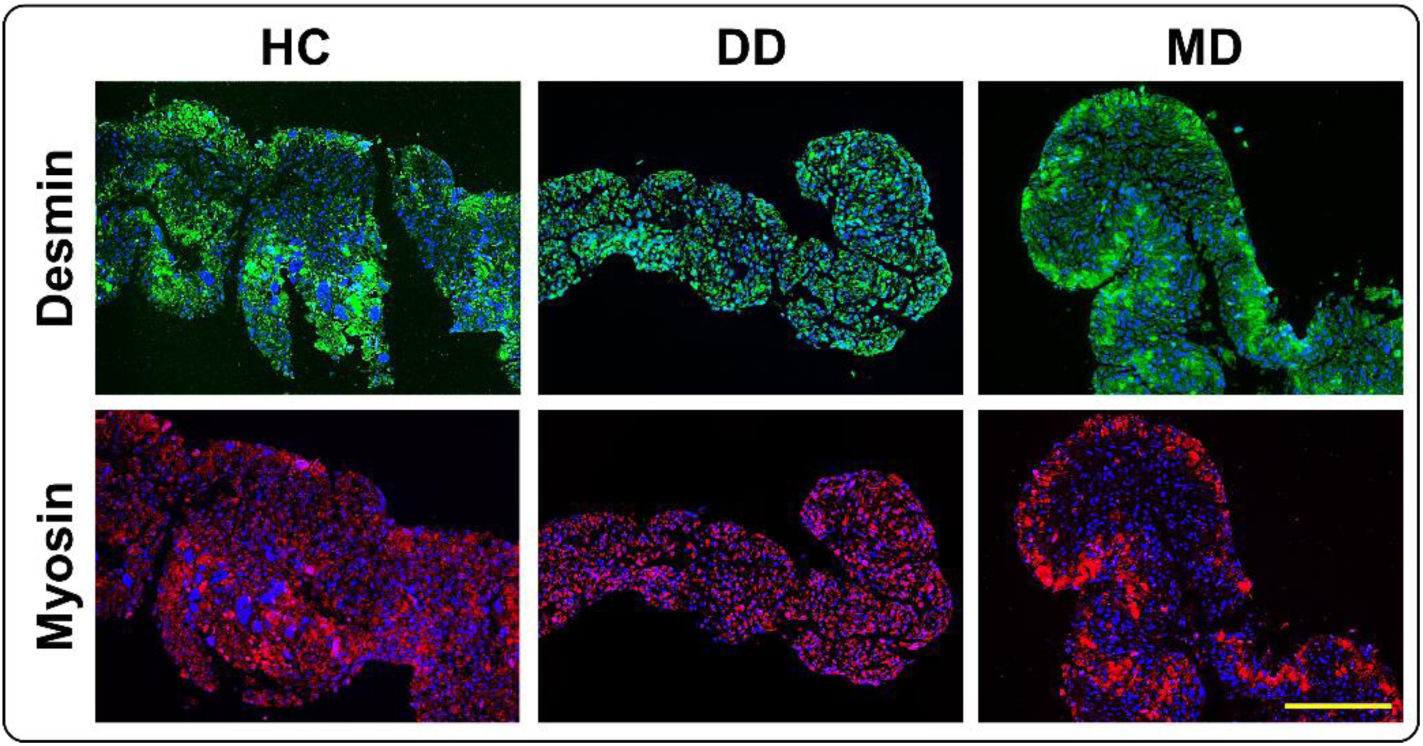
Evaluation of differentiation and maturation of 3D anchored muscle fibers using immunostaining for desmin and myosin heavy chain (MyHC). All three groups showed proper staining for both targets indicating proper differentiation of the cells in the 3D anchored fibers. This staining also showed the structural organization and maturity of muscle fibers within the constructs, using desmin as a marker of muscle integrity and myosin heavy chain as an indicator of sarcomere assembly and muscle function. Proper differentiation and maturation of cells into mature muscle fibers demonstrates the models’ effectiveness in replicating key aspects of muscle tissue architecture and development. Fluorescent microscopy was performed using a BioTek Lionheart automated microscope with suitable filters. Scale bar is 200 µm.

**Table 1.** Details of commercially available primary human myoblasts acquired from CookMyosite®.

| Ethnicity | Tissue of origin | Additional information |
| --- | --- | --- |
| Caucasian | Vastus Lateralis | Healthy Male, No diabetes, Tobacco and alcohol user, BMI 31, 29 years old |
| Caucasian | Vastus Lateralis | Male diagnosed with Duchenne muscular dystrophy (deletion in the Dystrophin gene (Exons 47-52) with muscle wasting), No Diabetes, No Tobacco or alcohol use, BMI 26, 20 years old |
| Caucasian | Vastus Lateralis | Female diagnosed with Myotonic dystrophy type 1 (Allele 1:23 CTG repeats in the DM-1 gene; Allele 2: 90-100 CTG repeats in the DM-1 gene), No Diabetes, No Tobacco or alcohol use, BMI 20, 18 years old |

In the immunostaining analysis of desmin and MyHC in 3D anchored muscle fibers, all three groups exhibited evidence of proper differentiation. In HC group, desmin and MyHC showed robust staining, consistent with the well-organized sarcomeric structure expected in healthy muscle tissue, underscoring successful myogenic differentiation and sarcomere assembly. Despite the altered morphological features in DD and MD groups, the presence of desmin and MyHC confirmed active myogenic processes and mature muscle fiber formation. Some variability in staining patterns between the healthy and diseased groups highlighted the specific pathological signatures of DD and MD. This validation underscored the 3D models’ ability to mimic not only healthy muscle physiology but also the specific states of muscular dystrophies, offering a versatile platform for investigating disease mechanisms and exploring potential treatments.

### Histological and immunostaining assessments of 3D anchored muscle fibers

To further delineate the morphological differences between 3D anchored muscle fibers of HC, DD, and MD groups, hematoxylin and eosin (H&E), Movat’s Pentachrome, and Masson’s Trichrome staining was used (**Figure 3a**). In HC group, the staining revealed densely packed, uniformly distributed myofibers, closely resembling the structural integrity typical of normal muscle tissue observed *in vivo*. By contrast, the DD and MD groups displayed distinct pathological characteristics consistent with their respective disease phenotypes. These included variability in fiber sizes, expanded interstitial spaces, and increased fibrosis, as indicated by greater collagen deposition observed through enhanced blue/green staining in Movat’s staining. These histological features are emblematic of the muscle degeneration typically observed in these disorders *in vivo* [28]. While the models faithfully replicated many critical aspects of dystrophic muscle pathology, enhancements in cellular architecture, such as improved cell density and inter-fiber connectivity, could further increase their resemblance to *in vivo* conditions. This improvement could be achieved by the incorporation of additional relevant stromal cell types, including fibroblasts and pericytes, as well as neuronal elements that are crucial in the pathophysiology of these diseases. Stromal cells can potentially be introduced through the use of patient specific iPSC-derived cells [29].

**Figure 3.**
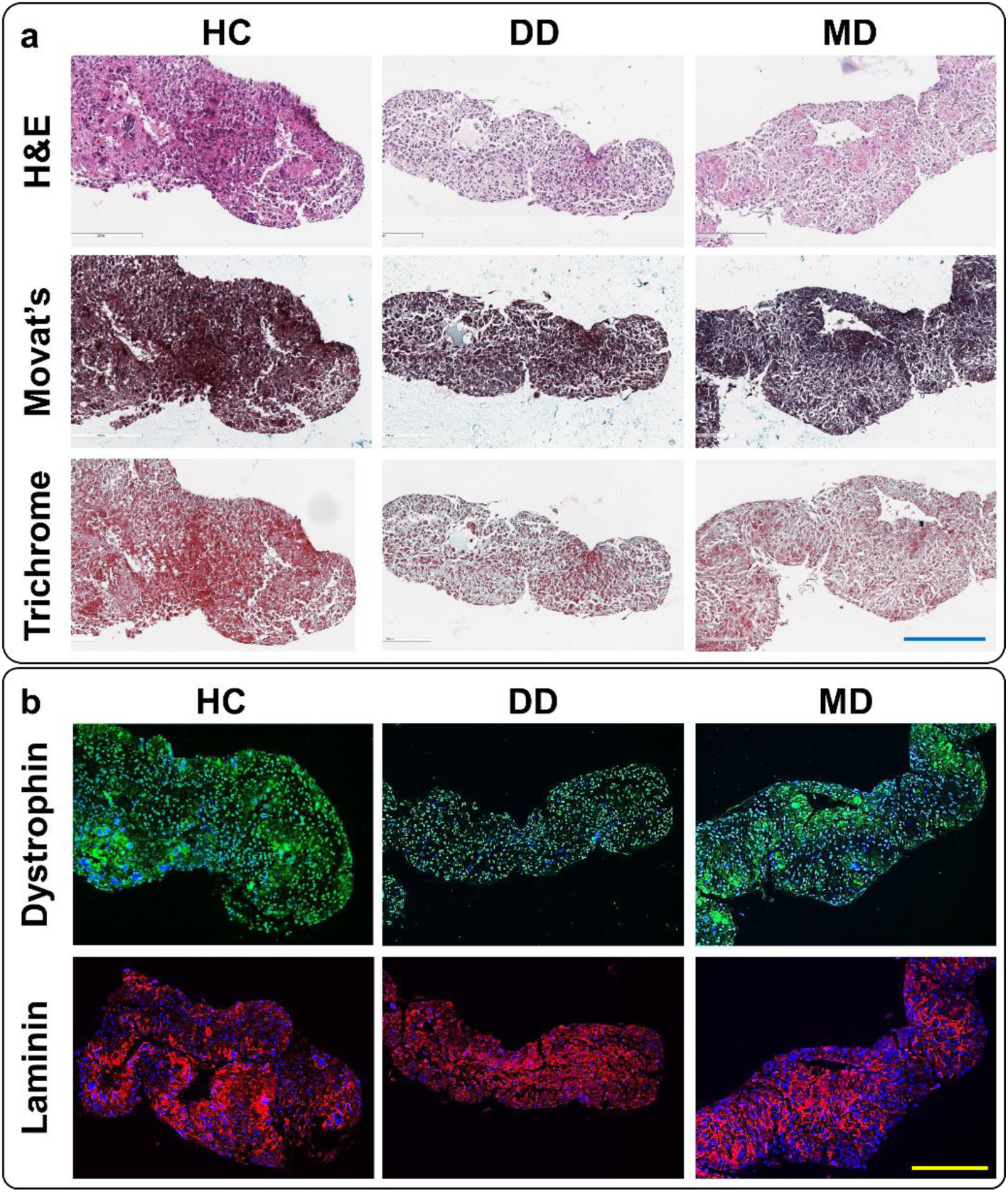
**a)** Histology and **b)** immunostaining of HC, DD, and MD 3D anchored muscle fibers. Hematoxylin and Eosin (H&E) stains nuclei blue and cytoplasm pink highlighting cellular and tissue structures. Movat’s Pentachrome differentially stains muscle fibers red, collagen blue/green, and elastic fibers dark brown and nuclei in black, illustrating the tissue organization and potential fibrosis. Masson’s Trichrome stains cytoplasm and muscle fibers in red/pink, nuclei black, and collagen blue, identifying muscle damage and fibrotic changes. Immunostaining targeted dystrophin, shown in green, indicated the integrity of the sarcolemma, and laminin, shown in red, highlighting the basement membrane structure. These markers provide insight into the structural and functional status of muscle tissue, with disruptions visible in diseased models. The bright field microscopy was performed with a Leica Aperio AT2 slide scanner at 40X magnification. Fluorescent microscopy was performed using a BioTek Lionheart automated microscope with suitable filters and 10X magnification. Scale bars are 200 µm.

Further evaluation of these 3D anchored muscle fibers was conducted through immunostaining analysis, specifically targeting dystrophin and laminin to assess their distribution and expression across the different conditions (**Figure 3b**). The HC group demonstrated consistent and continuous staining patterns for both dystrophin and laminin, indicative of intact basement membranes and sarcolemma integrity. In contrast, both DD and MD groups showed a marked reduction and discontinuity in dystrophin staining, mirroring the disrupted membrane integrity characteristic of these diseases, with pronounced effects observed in the DD group. Similarly, laminin staining, while still detectable, exhibited altered distribution patterns that likely reflected basement membrane remodeling, particularly in the DD group. These findings affirm the models ability to accurately reproduce the protein expression profiles characteristic of diseased muscle tissues, as well as the ultrastructural changes [30]. This provides a solid foundation for enhancing our understanding of muscular dystrophies and developing potential therapeutic approaches.

The differences in physical properties of the 3D anchored muscle fibers from different groups also reflected the differences in ECM and cellular organizations shown using histology and immunostaining. When the culture devices were subjected to shaking, the fibers showed varying responses due to their inherent structural differences. The HC group demonstrated robust mechanical integrity, maintaining a stable configuration with minimal deformation/swinging while hanging between the anchoring pillars, even to some degree bending the flexible pillars. In contrast, the DD group exhibited significant flexibility, with the fibers swinging extensively in response to even mild movements. Meanwhile, the MD group displayed more rigid formation, showing almost no movement upon shaking (**Supplementary Video 1**). These observations highlighted the distinct physical properties of the muscle fibers, aligning with the anticipated pathological characteristics of the respective muscular dystrophies and providing a functional demonstration of tissue phenotype under dynamic conditions. Similar variations in mechanical properties between healthy skeletal muscle tissues and muscles affected by different types of dystrophies have been documented *in vivo* [31–34].

### Differences in proteomics profile of 2D and 3D models

Proteomics analysis was conducted to compare 2D and 3D conditions within each healthy and diseased group, and to demonstrate the enhancements in 3D anchored fibers relative to *in vivo* conditions. The 2D conditions were prepared according to standard protocols. Culture was started with 5×10^5^ cells per condition in growth medium on day 1 in 6 well plates. By day 3, when confluence was nearly reached, the medium was switched to differentiation medium, which was refreshed bi-daily. On day 8, the cultures were washed with PBS, scraped, and the resulting cell pellets were frozen for subsequent analysis. Differentiation was verified by imaging and the presence of multinucleated myofibers (**Supplementary Figure 1**). Label-free proteomics was applied to two samples for each condition. Similarly, two samples of 3D anchored muscle fibers from each group were prepared as previously described, followed by proteomics analysis. Protein identification and quantification were carried out using Proteome Discoverer. Mass spectrometry data were then analyzed with MaxQuant software for peak identification, quantification, and normalization. Scaffold software was used for further validation, setting a 95% confidence threshold for protein identification and a minimum requirement of two unique peptides per protein to ensure the reliability of the proteomics data. Quantitative protein data were reported as normalized total precursor intensity, utilizing the label-free quantification approach to robustly measure protein abundance across samples. A comprehensive list of detected proteins (total of 3530 proteins) and their quantified expression levels is provided as **Supplementary File 1**.

In order to compare 2D and 3D conditions of each of the HC, DD, and MD groups, differential expression analysis was conducted using Perseus software, with first applying log2 transformation to the data followed by ANOVA testing (S0=0, FDR=0.05) over 250 randomizations for robustness. Results were visualized through volcano plots (**Figure 4**) generated by a custom Python script, identifying differentially expressed proteins under conditions of P-value < 0.01 and a minimum fold change of 10. Proteins were categorized and color-coded in the plots as upregulated (red), downregulated (blue), or non-significant (gray). The full list of up- and down-regulated proteins in each comparison is available as **Supplementary File 2**. Next, functional enrichment was performed by manually importing differentially expressed proteins in each case to g:Profiler (https://biit.cs.ut.ee/gprofiler/gost) against Gene Ontology (GO) (BP: Biological Processes, CC: Cellular Component, MF: Molecular Function). The full list of enriched pathways as well as other details including involved proteins in each pathway, full visual representation of enriched pathways, and statistical significances are included as **Supplementary File 3**.

**Figure 4.**
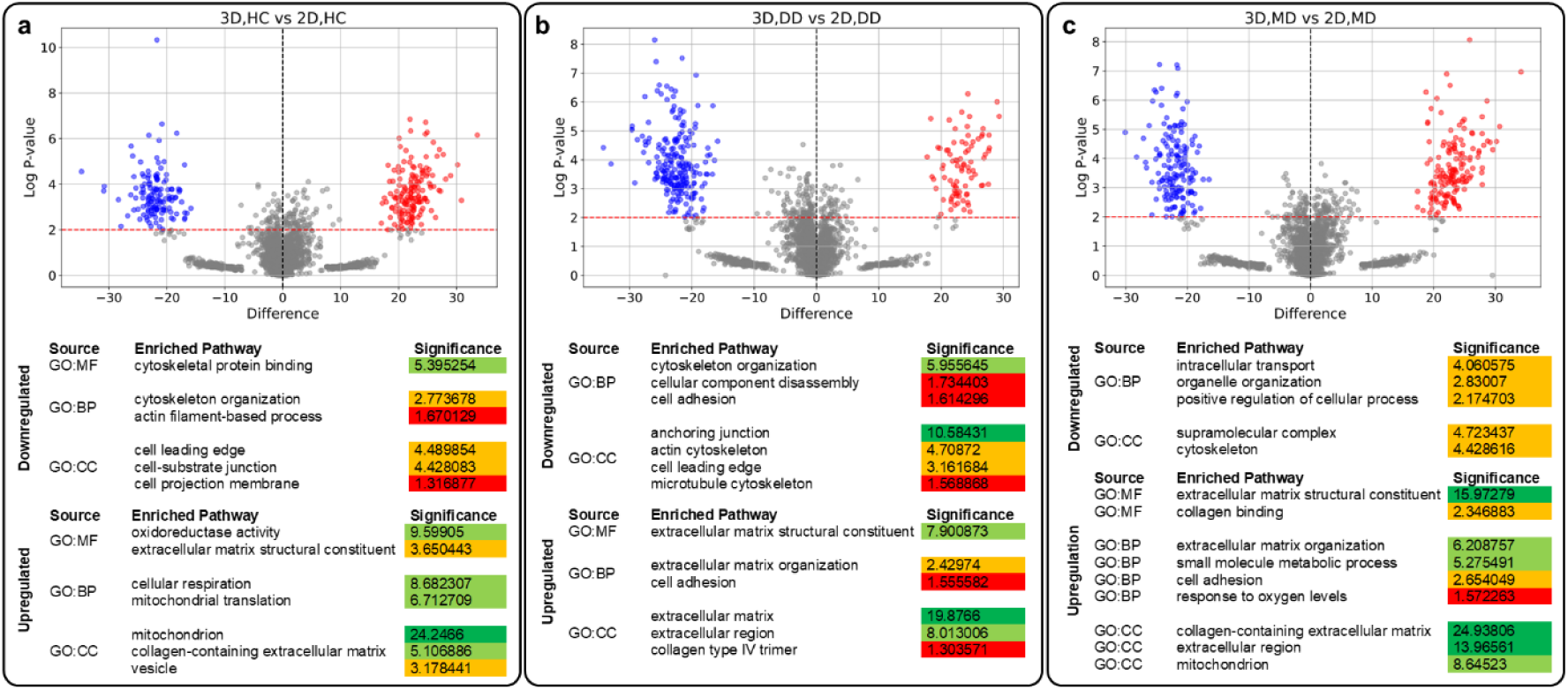
Differential expression analysis and enriched pathways when comparing 3D anchored fibers with 2D models for **a)** HC, **b)** DD, and **c)** MD groups. In all three comparisons, enriched pathways in up- and down-regulated proteins represent more *in vivo* like representations in 3D anchored fibers. Only some of the enriched pathways are shown here. Full list of enriched pathways is included in **Supplementary File 3**. Significance is defined as -log2 of adjusted P-value calculated by g:Profiler. Significance values are color coded with dark green showing values above 10, light green between 5 and 10, orange between 2 and 5, and red below 2.

When comparing 2D models of all three groups with their 3D anchored fibers (**Figure 4**), pathways related to the extracellular space, including ECM constituents and organization, were upregulated in the 3D anchored fibers, consistent with the formation of proper microstructures in 3D, as confirmed by histology and immunostaining. Similarly, in most instances, cell adhesion pathways, along with mitochondrion and metabolism pathways, were upregulated in all three 3D groups. This upregulation correlated with increased ECM content and 3D structure formation, as well as enhanced cell density and increased construct thicknesses changing mass transfer balances. At the same time, in all three groups, pathways related to cytoskeleton organization and components, particularly actin filament and microtubule were downregulated in 3D, indicating the formation of stable and mature constructs. In line with this, pathways related to cell morphology and intracellular organization, including cell projection membrane, cell leading edge, and other cellular processes, were also downregulated in 3D. This further highlighted the increased similarity of 3D anchored fibers to *in vivo* conditions, evidenced by the deposition of proper ECM and the formation of mature and stable constructs.

### Contrast in proteomics profile of diseased and healthy anchored fibers

When comparing 3D anchored fibers of dystrophic groups, DD and MD, with healthy HC group through differential expression analysis, meaningful *in vivo* like patterns emerged (**Figures 5a and b**). In both comparisons, pathways related to ECM and extracellular region were upregulated. In diseased human muscle, aberrant deposition of ECM and fibrosis happen to compensate for the loss of tissue integrity [35–37]. In these dystrophic anchored fibers, ECM components, such as Collagen alpha-1(VIII) chain (P27658), were upregulated, suggesting fibrotic tissue formation. This was consistent with the established roles of collagen type VIII in muscle fibrosis in humans [38, 39]. In DD group, downregulation of pathways related to muscle structure and its constituents, as well as muscle contraction and calcium ion bonding necessary for muscle function, were apparent. The observed downregulation of cell surface components, including cell adhesion molecules and junctions, also reflected the loss of components such as dystrophin in Duchenne dystrophy. Beta-1-syntrophin (Q13884) was significantly downregulated in DD group but not in MD. Dystrobrevin alpha (Q9Y4J8), a protein involved in the dystrophin-glycoprotein complex, along with Beta-1-syntrophin (Q13884), a component of the dystrophin complex, were also significantly downregulated. This further contributed to the destabilization of muscle tissue integrity. The role of syntrophin-dystrobrevin complex in human neuromuscular disorders has been described in detail before [40]. Components such as Tensin-3 (Q68CZ2), responsible for connecting ECM to cytoskeleton, as well as Integrin alpha-3 (P26006), mediating the adhesion of cells to the ECM, were downregulated in DD model similar to *in vivo*. Roles of Tensin and Integrin family members in skeletal muscle regeneration and function have been discussed before [41, 42]. The upregulation of Pappalysin-1 (Q13219), a secreted zinc-binding metalloproteinase, alongside the downregulation of Metalloproteinase inhibitor 3 (P35625), further verified the dysregulation of ECM in these dystrophic anchored fibers. This pattern mirrored the established roles of matrix metalloproteinases and their inhibitors in the inflammation and fibrosis of skeletal muscles [43]. When comparing MD group to HC, multiple MyHC isoforms (D6RJ91, P10916, Q92614) were downregulated, indicating damage to the muscle tissue fibers’ structure. Similar events are described for MyHC family members in different muscle dystrophies in both Duchenne [44] and Myotonic [45] dystrophies. Interestingly, the MD group showed fewer significantly upregulated or downregulated proteins when compared to the HC group, relative to the DD versus HC comparison. This reflected the more complex nature of Myotonic dystrophy, which involves multiple tissues beyond just skeletal muscle and showed the need for inclusion of other cell types and tissues in order to further increase the *in vivo* relevance of the model.

**Figure 5.**
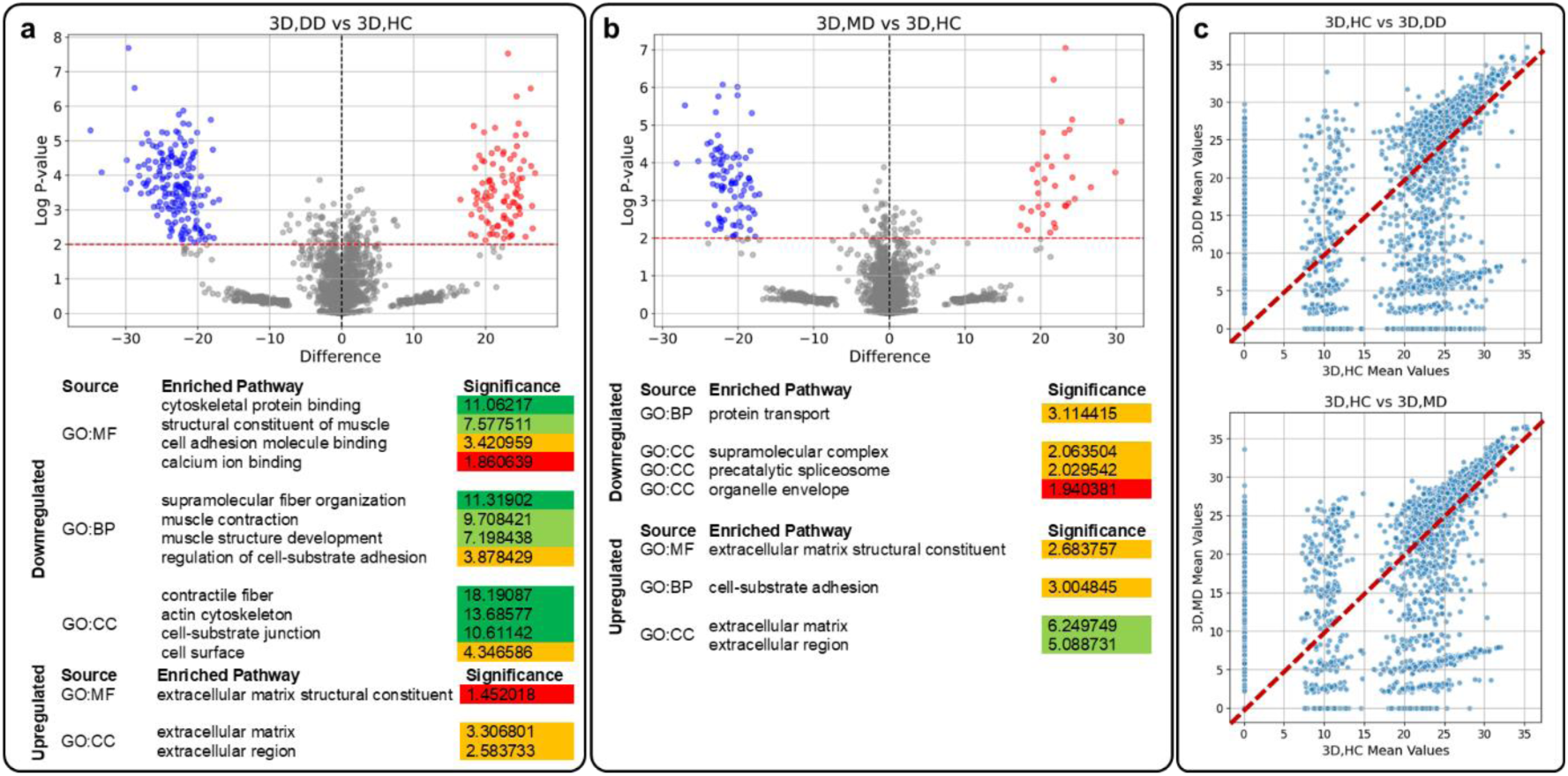
Differential expression analysis and enriched pathways comparing **a)** 3D anchored fibers of DD and HC and **b)** MD and HC groups. **c)** Comparison of overall expression patterns between healthy and diseased groups through scatter plots. In both comparisons, up- and down-regulated enriched pathways showed proper and *in vivo*-like recreation of differences between diseased and healthy phenotypes in 3D anchored fibers. Only some of the enriched pathways are shown here, and the full list is available as **Supplementary File 3**. Significance is defined as -log2 of adjusted P-value calculated by g:Profiler. Significance values are color coded with values above 10 shown in dark green, 5-10 in light green, 2-5 in orange, and below 2 in red.

The overall expression pattern of proteins for both DD and MD groups were compared with HC group through scatter plots (**Figure 5c**). In these plots, proteins with more similar expression levels in the two conditions being compared positioned closer to the diagonal. Conversely, proteins that appeared further from the diagonal had larger differences in their expression levels. In these comparisons, the majority of proteins were distant from the diagonal line, representing the gap between dystrophic and healthy groups. In both cases, two major clusters of proteins with vertical distributions, each containing smaller and closely positioned sub-clusters, were detectable. Notably, some proteins were only expressed (showing nonzero expression values) in just one of the conditions being compared, further widening the disparity between the two groups and highlighting the significant differences between dystrophic and healthy conditions.

### Assessment of drug treatment effect on muscular dystrophy models

DD and MD 3D anchored muscle fibers were treated with creatine (1 mg/mL) and mexiletine hydrochloride (2 mg/mL), respectively, to assess whether drug treatment could improve their phenotype and structure to more closely resemble healthy conditions. Treatment was started as soon as the 3D fibers were formed on day 21, 24 hours after scraping. The desired amount of the drug was added to the differentiation medium, and treatment was continued until day 24 when the constructs were fully formed and matured. Creatine has been used in subjects suffering from Duchenne dystrophy [46] with patients experiencing improved muscle performance during its supplementation [47]. Similarly, mexiletine hydrochloride is used with Myotonic dystrophy patients [48], and hand grip myotonia improvements have been reported in these cases [49]. However, no study has evaluated the effect of either of these components on *in vitro* models of skeletal muscle tissue to date. At first glance, treatment with mexiletine showed an obvious decrease in the stiffness of the MD anchored fibers (**Supplementary Video 2**). To further study the effect of drug treatment on these anchored 3D fibers, proteomics analysis was performed as well, and drug treated conditions were compared with both their untreated counterparts and the HC group.

Volcano plots showed a high number of proteins down- or up-regulated in both cases (**Figure 6**) with many different pathways enriched in each case (**Supplementary File 3**). In both DD and MD groups, comparing drug treated conditions with their untreated counterparts using scatter plots showed formation of three major clusters positioned in proximity og the diagonal. Each of these clusters contained multiple small and closely positioned sub-clusters, some diverging from the diagonal line. When drug-treated conditions were compared with the HC group, six distinct clusters emerged, with two on the diagonal and four positioned away from it. The majority of the proteins were in cluster number 4, and some in cluster 3, both positioned on the diagonal, while the other clusters, situated away from the diagonal, indicated a lack of positive effect from the drug treatment in shifting the protein expression pattern to more closely resemble that of the HC group for these clusters. Next, protein-protein interaction networks were generated for proteins in cluster 4 using STRING tool (https://string-db.org/) for both DD and MD groups treated with drugs. The majority of the proteins in this cluster were associated with the Extracellular Region, Cell Junction, or Mitochondrion, according to the cellular composition gene ontology analysis (**Supplementary Figures 2 and 3**). This suggested a high similarity of proteins particularly crucial in both diseases when comparing drug-treated diseased models and the HC group, indicating a shift toward a healthier phenotype. The presence of clusters away from the diagonal in these comparisons reflected the complex nature of these diseases and potentially the need for more sophisticated drug treatment regimens, where multiple drug types are used to treat different aspects of each disease.

**Figure 6.**
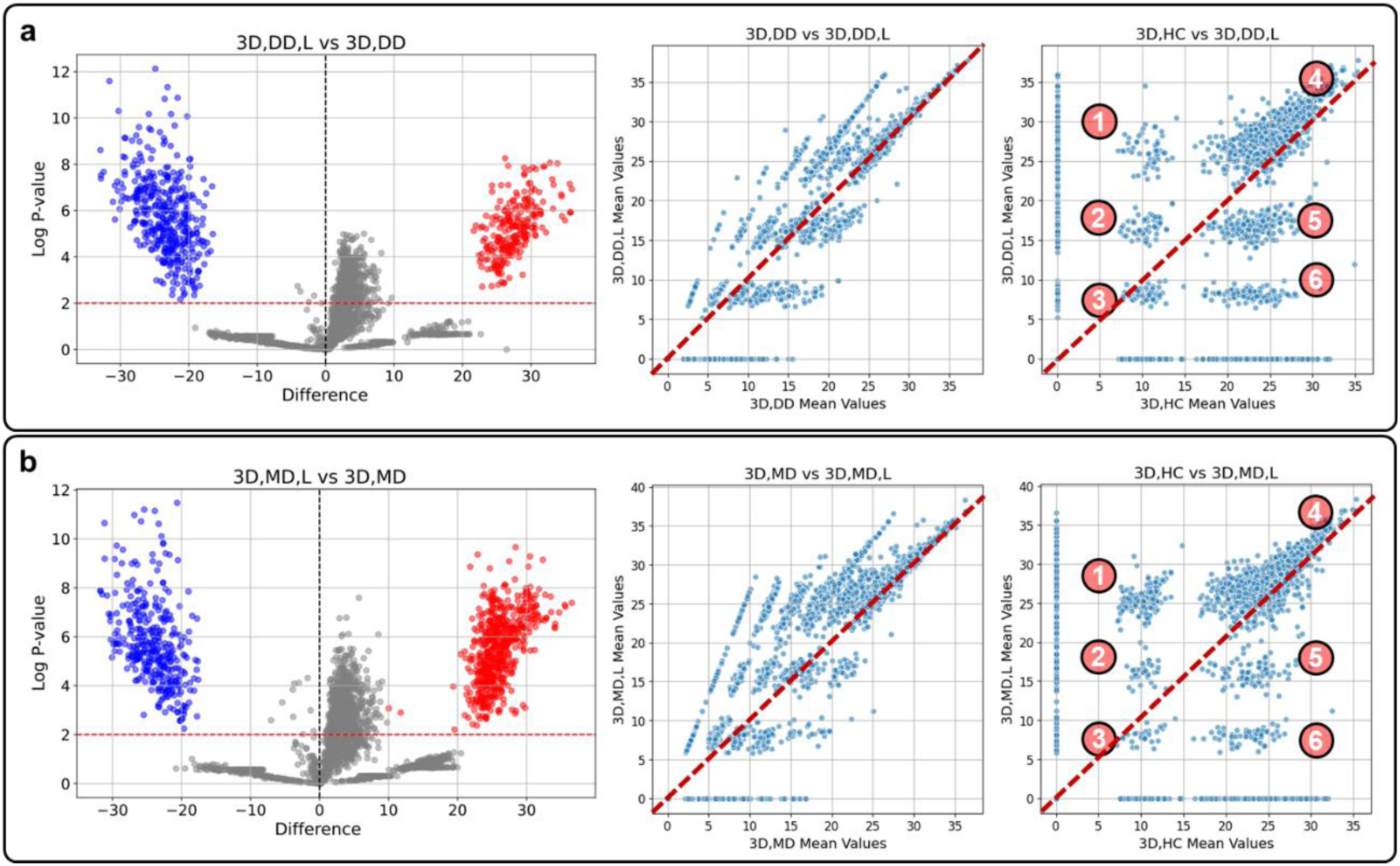
Differential expression analysis and scatter plots investigating 3D anchored fibers of **a)** DD group treated with 1 mg/mL creatine and **b)** MD group treated with 2 mg/mL mexiletine hydrochloride. While differential expression analysis revealed a significant number of proteins as up- or down-regulated in each case, comparing drug-treated conditions with their untreated counterparts using scatter plots showed some similarities between the two. At the same time, when drug treated DD and MD groups were compared with HC group, 6 distinct clusters were formed, with a majority of the proteins showing high similarity between the two conditions (clusters 3 and 4), some showing small changes (clusters 2 and 5), and the rest showing bigger differences (clusters 1 and 6). L stands for lower concentration drug treatment in each case.

The effect of increasing the drug concentration in each treatment was also studied (**Supplementary Figure 4**). Both volcano plots and scatter plots showed the same pattern in changing protein expression levels when a higher concentration of each of the drugs were used, but the degree to which these changes happened was different. When lower and higher drug concentration conditions were compared with each other through scatter plots (**Supplementary Figure 5a**), much bigger changes were revealed, with 9 distinct clusters forming in each case, three of which on the diagonal and the rest away from it, showing the importance and effect of drug concentration on treating these conditions. Interestingly, despite the inherent differences between Duchenne and Myotonic dystrophies, the overall patterns of cluster formation in these comparisons had stark similarities.

To better compare overall differences in protein expressions in different diseased and drug treated 3D anchored muscle fibers with HC group, UpSet plots were used to depict the general increases and decreases in expression values of proteins (**Figure 7a**). First, log10 transformation was performed for protein expression levels to stabilize variance and normalize the distribution, addressing potential issues of heterogeneity common in proteomic data. To ensure the comparative analyses reflected true biological variations rather than technical biases, quantile normalization was applied to align the distribution of protein expression values across all samples. Then, the mean value was calculated for each protein across all replicates. Considering 3D anchored fibers of HC group as control, Z-scores were calculated for each protein in other conditions relative to this control, providing a measure of deviation in units of standard deviation, making it possible to identify significant changes in expression levels. A threshold Z-score of ±1, roughly equivalent to the 84th percentile, was selected to detect a broad range of expression changes, capturing more subtle variations. This approach helped in identifying proteins with generally higher or lower expressions compared to the control, moving beyond a focus solely on statistically significant changes. Data visualization was conducted using UpSet plots to display the distribution and overlap of proteins with altered expression across conditions. Increased and decreased expression levels were highlighted with red and blue colors, respectively. Notably, some groups shared a significant overlap in protein expression profiles, suggesting common regulatory mechanisms or similar responses to experimental treatments. While the use of a Z-score threshold of 1 allowed for the inclusion of subtle expression changes, the need for cautious interpretation should be considered, as these changes encompass a wider range of biological variability.

**Figure 7.**
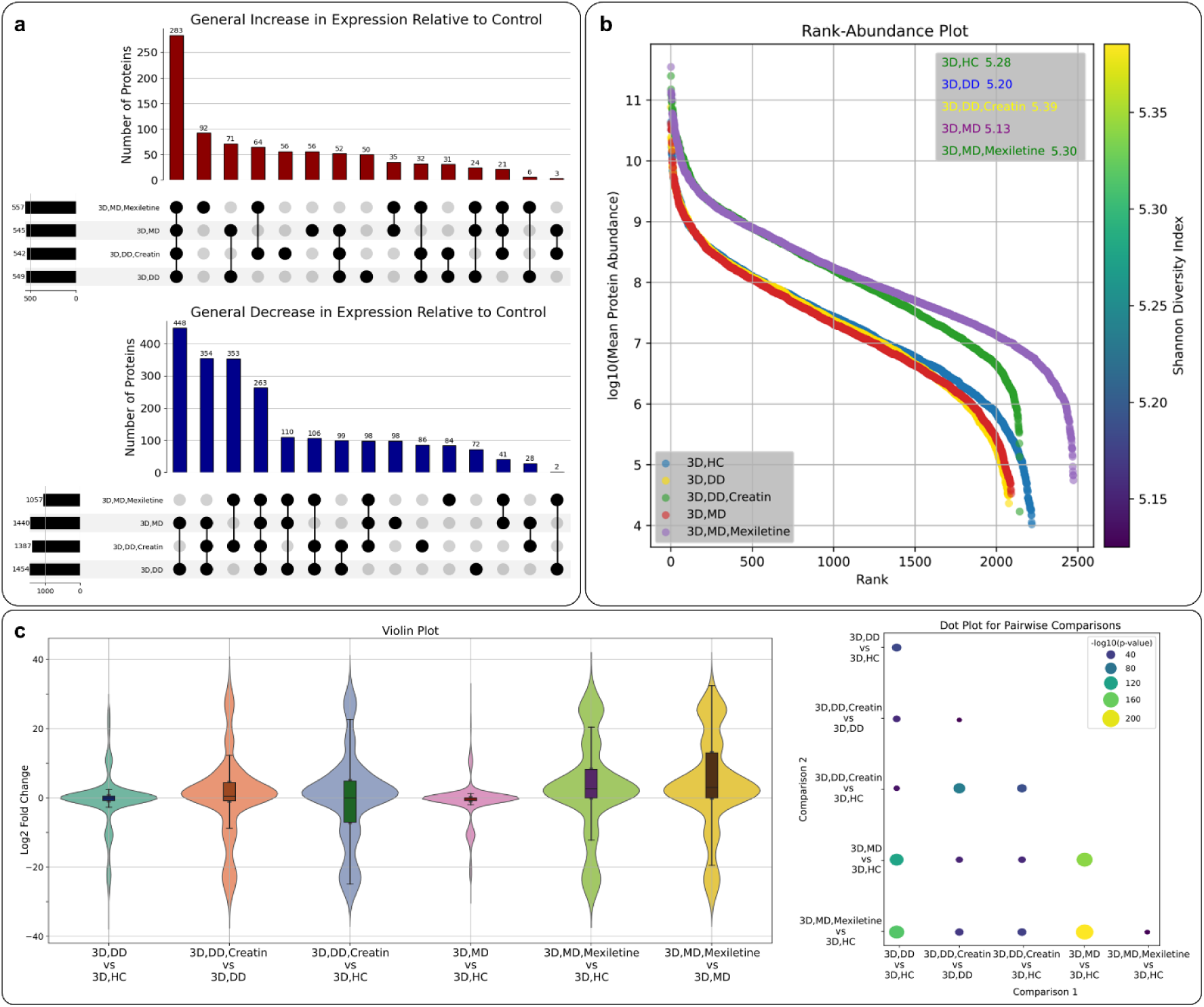
**a)** UpSet plots representing the number of proteins with increased or decreased expressions in diseased anchored muscle fibers and their drug treated counterparts compared to HC group, **b)** Rank-Abundance plots showing abundance ranges of the detected proteins and the Shannon Diversity Index showing diversity and richness of proteins in each group, **c)** Violin plots showing the distribution of fold changes in protein expressions when different groups were compared and dot plot showing the statistical differences between comparisons. In UpSet plots, each horizontal bar on the left side of the plot represents the total number of proteins that meet the criteria in a single group. Each vertical bar above shows the intersection size, i.e. the number of proteins commonly increased or decreased across the groups connected by dots in the matrix below. In the violin plots, the width at a given y-axis value indicates the density of data points (i.e., the number of proteins with that particular fold change) and the vertical range of the plot on the y-axis shows the spread of fold changes. A taller plot indicates a wider range of changes in protein expressions. Symmetry about the central line (log2 fold change of 0) means balanced increases and decreases in expression, while asymmetry indicates a bias towards either increased or decreased expression.

These groups were also compared with each other using a Rank-Abundance plot (**Figure 7b**). The log10 transformation was first performed for proteins, allowing for the identification of subtle differences in the higher and lower abundance ranges, which are often overlooked. These visualizations provided an intuitive mean to assess and compare the protein landscapes across different experimental conditions, enhancing understanding of the proteomic complexities associated with each condition. Interestingly, DD and MD 3D anchored muscle fibers had similar profiles here, but the drug treatments changed their profiles, with both drug treated DD and MD profiles being extended on x-axis, showing higher numbers of detected proteins. Creatin treated DD profile on x-axis was similar to HC condition as well. Both drug treated groups also showed a shift upwards on the y-axis, indicating an increase in overall expression levels of detected proteins. To better quantify the diversity of protein expression patterns across different conditions, the Shannon Diversity Index was calculated. This index accounts for both the richness, total number of different proteins, and evenness, similarity in expression values, of the proteomics profile in each case. A higher Shannon Diversity Index indicates greater diversity. For both the DD and MD cases, drug treatment increased the Shannon Index in both instances, making them closer to HC group, suggesting a potential therapeutic benefit through enhanced protein diversity. These differences between different groups were not apparent in other analysis and showed the importance of evaluating the effect of drug treatment using multiple analysis types. When groups of low and high drug concentrations were compared using Rank-Abundance plots and the Shannon indices (**Supplementary Figure 5b**), the overall change in patterns was similar in each case with some minor differences, serving as further evidence of the importance of optimizing drug concentrations.

To further illustrate the differences between various groups and the effects of drug treatments in each disease, the distribution of fold changes in protein expressions were visualized using Violin plots for different comparisons (**Figure 7c**). The plots were generated using a custom Python script and the “difference” values calculated by Perseus as part of the differential expression analysis performed on the log2 transformed expression intensities. This visualization highlighted the differences in the distribution of protein expression changes in each comparison. Based on these results, both DD and MD groups exhibited lower protein abundances and significant changes in protein expressions compared to the HC group. Creatine showed low impact on the DD group in terms of protein abundance and expression compared to both the HC and DD groups. Mexiletine hydrochloride had a more pronounced effect on the MD condition, increasing protein abundance and shifting protein expression levels closer to the HC group. In their respective comparisons, both drug-treated groups displayed taller plots, indicating a wider range of protein expression changes. This suggested that the drug treatments caused significant variations in protein levels. The drug-treated groups also showed wider plots, indicating a higher density of proteins experiencing similar fold changes. These observations were consistent with the formation of multiple smaller clusters when the data was visualized using scatter plots. The drug-treated groups exhibited a small upward shift, suggesting an overall increase in protein expression intensities in both comparisons. This is in line with the upward shift observed in the Rank-Abundance plots and higher Shannon Diversity Indices. These Violin plots further complemented the insights provided by other analysis presented here.

To ensure the observed differences in protein expression were not only visually apparent but also statistically significant, a rigorous statistical approach was implemented. For each pair of comparisons, the script conducted a series of statistical tests to determine statistical significance. The Shapiro-Wilk test checked for normality, and the Levene’s test assessed variance equality. Based on these assessments, an independent t-test (used if both comparisons passed the Shapiro-Wilk test for normality and Levene’s test showed equal variances), Welch’s t-test (used if both comparisons passed the Shapiro-Wilk test for normality, but Levene’s test showed unequal variances), or Mann-Whitney U test (used if either comparisons failed the Shapiro-Wilk test for normality, regardless of the results from Levene’s test) was applied as appropriate. The Benjamini-Hochberg procedure adjusted P-values to control the false discovery rate. The analysis results, including test types, statistics, and adjusted P-values are included as **Supplementary File 5** and visualized using a dot plot (**Figure 7c**). The combination of violin plot and statistical tests provided a comprehensive view of the differential expression data, highlighting both the overall distribution and the specific significant differences between the comparisons, enhancing the robustness and reliability of the findings presented here.

Since routinely used 2D culture models fail to properly recapitulate the structure and function of skeletal muscle tissue for healthy and diseased conditions, the development of functional and translatable 3D models has become a priority [50, 51]. Despite significant advancements in recent years, majority of the 3D models, constructed using techniques such as bioprinting, micromolding, micropatterning, and scaffold-based methods such as electrospinning, still depend heavily on biomaterials such as collagen, Matrigel®, decellularized ECM, alginate, gelatin, and gelatin methacrylate, among others. These materials often lack the specific recognition moieties crucial for proper cell adhesion, migration, and function, or they are not customized for specific tissues and diseases, leading to their failure to induce appropriate cellular phenotypes. Most of these studies continue to rely on the mouse immortalized C2C12 cell line, which falls short in replicating many human-like features, not to mention the specific phenotypes associated with different diseases [15]. The use of patient-derived primary muscle cells [12, 52] or patient-derived iPSCs [11, 53] has become more prevalent recently. However, very few studies have used scaffold- and biomaterial-free platforms that incorporate pillars or micropillars to manage the physical properties of developing tissues [54], and none have used cell sheets or appropriately addressed the significance of the ECM in their frameworks. Other scaffold-free but anchor-less modeling techniques such as myospheres made using hanging drop technique or low attachment surfaces [15, 51, 55] show reduced mechanical stability, are limited in size and complexity/maturation, and suffer from lack of reproducibility.

The anchored cell sheet engineering platform utilized in this study, in conjunction with patient-specific primary cells, successfully addressed previous limitations and faithfully recreated *in vivo* skeletal muscle conditions. This platform shows promise for mechanistic studies and drug discovery, representing a significant potential step toward linking laboratory research with clinical applications. By using patient- specific cells to recreate disease-specific ECM and tissue architecture, this method enables precise evaluations of drug efficacy and safety in a context that closely resembles human muscle physiology, substantially reducing the time and resources required to assess potential treatments prior to clinical trials. Additionally, the capability to model various stages of disease progression offers essential insights into the optimal timing and dosing of therapeutic interventions, enhancing the likelihood of clinical success. In addition, the potential for this platform in regenerative medicine is also considerable. For instance, it could be used to treat conditions such as volumetric muscle loss by implanting scaffold-free, fully mature muscle fibers derived from either patient-specific stem cells or allogeneic cells.

## 3. Methods

### Anchored cell sheet biofabrication process

A previously developed scaffold-free biofabrication platform based on the cell sheet engineering concept, called anchored cell sheet engineering [24], was used to develop 3D anchored fibers of skeletal muscle tissue for *in vitro* modeling. Briefly, culture devices were fabricated using a 3D printed acrylonitrile butadiene styrene (ABS) master mold with inverse design of the culture device and casting soft Ecoflex™ 00-30 pillars as anchors, PDMS as the base of the culture device for adherent cell culture, and a blend of PDMS and Mold Star™ 15 for the culture device walls to contain the medium. After the resins were cured, the culture device was peeled off from the master mold and the 3D printing surface textures composed of parallel patterns with shallow grooves were replicated on the PDMS base for inducing cell alignment and higher ECM production caused by 2.5D nature of the culture. The hydrophilicity of the PDMS base membrane was adjusted through treatment with a 50 mg/mL aqueous solution of tannic acid (Sigma-Aldrich, 403040) in deionized water, for a duration of 72 hr, followed by multiple washing rounds with deionized water to eliminate tannic acid residues and then autoclaving. Culture devices were then treated with a 50 µg/mL vitronectin (Gibco™, A14700) solution in PBS for one hour at room temperature and cell culture was started afterwards.

For culture purposes, each culture device was secured on a 3D printed polyethylene terephthalate glycol (PETG) support sterilized with 70% ethanol and were maintained in a sterile petri dish. On day one, culture was started by adding 5×10^5^ of each primary myoblasts in 3 mL of growth medium to achieve 80% confluency by the next day. On day 3, with cells at near 100% confluency, culture was switched to cells’ differentiation medium. This medium was refreshed once every three days, while additional 5×10^5^ cells were introduced to each device on days 4, 7, 10, 13, 16, and 19. By day 20, a multi-layered cell and ECM growth loosely attached to the surface of the membrane had formed. This layer was released from the membrane using a sterile 16 mm blade scraper (Sigma-Aldrich, C6106) by gently scraping the layer from four edges of the culture device. After releasing the cell sheet from the base membrane and losing its anchorage, the presence of the pillars was sensed by the cell sheets and over the course of a 24-hour process, cells remodeled the sheet to form a 3D fiber suspended between two pillars acting as new anchors for the now 3D constructs. Another 72 hours in culture was allowed to allow fibers to maturate.

### Primary cells and culture conditions

Commercially available primary myoblasts, healthy, Duchenne dystrophy, and Myotonic dystrophy type 1, were acquired from CookMyosite® (**Table 1**). Cells were grown in CookMyosite’s Myotonic™ growth medium (Basal medium (MB-2222-A-083) with growth supplement (MS-3333-039)) in cell culture treated 10 cm petri dishes up to 70% confluency. After dissociating the cells using TrypLE™ Express Enzyme (1X) (Gibco™, 12604013) and replating them in the culture devices, differentiation was started at about 100% confluency using CookMyosite’s Myotonic™ differentiation medium (MD-5555-2-023). Differentiation of the cells was confirmed visually using bright field microscopy.

### Drug treatment

Response of the 3D anchored muscle fibers from DD and MD groups to drug treatment was evaluated by treating them with 1 mg/mL creatine (SigmaAldrich, C0780), and 2 mg/mL mexiletine hydrochloride (SigmaAldrich, M2727), respectively. Treatment was started on day 21, 24 hours after scraping had done and the 3D tissue fibers had formed. The desired amount of the drug was added to the differentiation medium, and treatment continued for 3 days covering the rest of the maturation process. At the end of the process, 3D anchored fibers were washed with PBS and frozen down for further proteomics analysis.

### Sample processing, histology, and immunostaining

Biofabricated 3D anchored muscle fibers were sectioned using a standard microtome. Sections were then subjected to immunostaining to investigate the protein expression of myosin (Abcam, ab37484), laminin (Abcam, ab11575), desmin (Abcam, ab32362), and dystrophin (Abcam, ab275391). Primary antibodies were used at a dilution of 1:500 and for the detection of primary antibodies, secondary antibodies were employed at a dilution of 1:1000. Alexa Fluor 546 goat anti-mouse (Abcam, ab150116) was used for myosin, and Alexa Fluor 594 goat anti-rabbit (Abcam, ab150080) was utilized for laminin, desmin, and dystrophin. In addition to immunostaining, histological staining was performed to assess the structural integrity and morphology of the biofabricated muscle tissues. The staining methods included Hematoxylin and Eosin (H&E), Movat’s Pentachrome, and Masson’s Trichrome. Staining was executed following standard procedures to highlight different tissue components and fibrous structures. Fluorescent and phase contrast microscopy was done using a BioTek Lionheart automated microscope with suitable filters and 10X magnification. Images were captured under consistent lighting and magnification to ensure accurate representation of staining patterns and protein localization. The slides were scanned with Leica Aperio AT2 slide scanner at a 40X magnification.

### Proteomics analysis

Proteomic analyses were conducted on 2D cultures and 3D anchored muscle fibers, in each case in replicates. Protein extraction was achieved using TONG Lysis buffer supplemented with 10 mM TCEP and 40 mM Chloroacetamide, followed by sonication (15 sec, 4 Watt) and chloroform/methanol protein precipitation. Protein digestion was performed with 3.0 µg of Trypsin per sample, with enzyme-to-substrate ratios ranging from 1:50 to 1:200, at 37°C overnight. Post-digestion, samples were acidified with 1% trifluoroacetic acid, concentrated using a Speedvac, and desalted using Millipore C18 ZipTips. Following lyophilization, peptides were resuspended in 2% acetonitrile with 0.1% formic acid, and analyzed using a Thermo Scientific Orbitrap Fusion Lumos mass spectrometer coupled to an Evosep One LC system. Chromatographic separation was achieved on a Dr Maisch C18 AQ column (1.9µm beads, 150µm ID, 15cm long) maintained at 60°C using a binary gradient of 0.1% formic acid in water and acetonitrile. Protein identification and quantification were performed using Proteome Discoverer version 2.5.0.400, searching against the Uniprot database with specific search parameters, including a parent mass error tolerance of 50 ppm and a fragment mass error tolerance of 0.6 Da. The analysis allowed for up to three missed cleavages and included several modifications such as carbamidomethylation, oxidation, deamidation, and acetylation.

Following mass spectrometry, the raw data were processed using MaxQuant software to perform peak identification, quantification, and normalization, employing accepted parameters to ensure rigorous analysis. The resulting output files were then imported into Scaffold software to further validate the protein identifications and quantifications. Scaffold was configured to apply a 95% confidence threshold for protein identification and a minimum of two peptides per protein, each meeting a 95% confidence threshold for peptide identification. This stringent filtering was crucial for enhancing the reliability of the protein data. Quantitative protein data were reported as normalized total precursor intensity, leveraging the label-free quantification approach to provide a robust measure of protein abundance across samples. For each of the healthy and diseased groups, in 2D and 3D conditions, two replicates were used while for drug treatment experiments, three replicates were considered for each condition.

### Differential expression analysis

Differential expression analysis was performed using Perseus software. Initially, quantitative protein data as normalized total precursor intensity values were imported to Perseus and categorical annotation rows were added to accurately label replicates, facilitating correct treatment grouping. Protein intensities, plus 1 to remove 0 values, underwent log2 transformation to normalize the data distribution, essential for subsequent statistical evaluations. ANOVA was conducted for multiple sample testing, setting the *S*0 parameter to 0 to reduce the influence of large fold changes on small mean differences, and applying a false discovery rate (FDR) of 0.05 to control error probability. The analysis included 250 randomizations to verify the robustness of the findings. After completing the statistical analysis in Perseus, the data were exported and analyzed further using a Python script to generate volcano plots. The script generated plots that visually illustrated the differential expression between conditions using computed values by Perseus through t-tests and demonstrating the -Log(P-value) on the x-axis and Difference values (log2 fold changes of proteins) on the y-axis. Differentially expressed proteins were identified under stringent criteria of P-value < 0.01 and a minimum fold change of 10. Proteins were categorized as upregulated (red), downregulated (blue), or non-significant (gray) in the plots. To facilitate visual analysis a dashed lines was added to the plots at the P-value threshold. Direct comparison of different conditions was performed through scatter plots, another Python script utilized the log2 transformed values for each condition. Each protein was represented by a dot on the plot, with its position indicating the mean log2 transformed values under the two conditions being compared.

### Pathway enrichment analysis

Differentially expressed proteins were analyzed for functional enrichment using g:Profiler (https://biit.cs.ut.ee/gprofiler/gost). Default settings were used, including g:SCS for multiple testing correction with a significance threshold of 0.05. Ambiguous IDs were ignored to ensure data clarity. Enrichment analyses were performed against Gene Ontology (GO) (BP: Biological Processes, CC: Cellular Component, MF: Molecular Function). Results were visualized and downloaded as both CSV and PNG files for further analysis.

### Comparative analysis of protein expressions across groups

The overall differences in protein expressions in different diseased and drug-treated 3D anchored fibers were compared to the 3D HC group using UpSet plots. The custom Python script followed several key steps to ensure accurate and meaningful comparisons. First, the protein expression levels were log10 transformed to stabilize the variance and mitigate the impact of extreme values. This transformation was followed by quantile normalization to ensure the distributions of protein expression levels across different conditions were comparable. For each condition, the mean expression value for each protein across these replicates was calculated. This provided a single expression value per protein per condition, facilitating straightforward comparisons. The 3D HC group was used as the control, serving as the baseline for comparative analysis. Z-scores were calculated for each protein in the other conditions relative to the control and a threshold Z-score of ±1 was selected to capture a broad range of expression changes, including subtle variations. UpSet plots were then generated to display the distribution and overlap of proteins with altered expression across conditions. Increased and decreased expression levels were highlighted with red and blue colors, respectively.

The overall differences in protein expressions in different diseased and drug treated 3D anchored fibers as well as the healthy group were compared using Rank-abundance plots. A custom Python script was used to calculate each protein’s mean abundance for the respective condition’s replicates and then log10 transformed to stabilize variance and improve interpretability. Proteins were ranked based on their median abundance, allowing for a comparative analysis across conditions. Distinct colors were assigned to each condition to facilitate clear differentiation in the visual representation. To assess the diversity of protein expression patterns across different conditions, the Shannon Diversity Index was calculated for each condition. This index provided a quantitative measure of diversity, accounting for both the abundance and evenness of protein expression levels. To calculate the Shannon Diversity Index, the proportion (*pi*) of each protein in a condition was first determined by dividing the protein’s expression intensity by the total expression intensity of all proteins in that condition. These proportion values were then used to calculate the Shannon Diversity Index using the formula (*H*=-∑^s^*i*=1*pi.*×log2(*pi*)) where s is the total number of proteins with non-zero abundance. A heatmap representation of the Shannon Diversity Index was generated to provide a comprehensive overview of the protein expression landscape.

The distribution of fold changes in protein expressions across different groups and comparisons (e.g., diseased vs. healthy, drug-treated vs. untreated, drug treated vs. healthy) was visualized using a custom Python script. The “difference” values calculated by Perseus, as part of the differential expression analysis on log2 transformed expression intensities, were used as inputs. A violin plot was then generated to visualize the distribution and overlap of protein expression changes across the different comparisons. To complement the visual analysis, statistical tests were performed to assess the significance of differences between the groups and comparisons. For each pair of comparisons, the script carried out multiple steps. First, the Shapiro-Wilk test was used to determine if the distribution of fold changes for each comparison was normally distributed, with a P-value greater than 0.05 indicating normality. Next, the Levene’s test assessed the equality of variances between the two groups, where a P-value greater than 0.05 indicated equal variances. Based on the results of these tests, the appropriate statistical test was selected. If both groups passed the normality test and had equal variances, an independent t-test was used. If the groups had unequal variances but were normally distributed, Welch’s t-test was applied. If one or both groups did not pass the normality test, the Mann-Whitney U test, a non-parametric test, was used. To account for multiple pairwise comparisons, the Benjamini-Hochberg procedure was applied to adjust the P-values and control the FDR. To do so, the P-values were sorted in ascending order and adjusted P-values were calculated using the formula *p_adj_*=*p_i_*×*n*/*i*, where *p_i_* is the P-value, *n* is the total number of tests, and *i* is the rank. Adjusted P-values were compared to the specified FDR threshold (0.05), and if they were below this threshold, it indicated a statistically significant difference. The results of the statistical analysis were compiled into a summary table, detailing the type of test used, test statistics, and both raw and adjusted P-values for each comparison. These results were also visualized using a dot plot.

## 4. Conclusions

This study showcased the effectiveness of a novel scaffold-free anchored cell sheet engineering platform in developing physiologically relevant, patient-specific *in vitro* models of dystrophic skeletal muscle. Through immunostaining for desmin and MyHC, full differentiation of cells within the 3D constructs was confirmed. These models not only exhibited structural integrity but also showed functional characteristics similar to native tissues, as evidenced by the proper distribution of key muscle proteins such as dystrophin and laminin. Histological analyses, including H&E, Movat’s Pentachrome, and Mason’s Trichrome staining, along with immunostaining assessments, demonstrated that these 3D models effectively replicated the pathological features of muscle dystrophies. These features included disrupted dystrophin distribution, which was compensated by excessive ECM deposition by the cells, highlighting the models’ ability to reproduce disease-specific cellular environments. Proteomics analyses further confirmed the superiority of the 3D models over traditional 2D cultures. These analyses showed that the 3D models more accurately reflected *in vivo* conditions, with differential expression analysis coupled with statistical evaluations revealing distinct proteomic profiles between diseased and healthy models. These differences underscored the accuracy and *in vivo* likeness of the approach. Additionally, with the use of pathway enrichment analysis, relevant biological pathways affected by the disease conditions were identified. Treatment of diseased models with clinically effective drugs, specifically creatine for Duchenne muscular dystrophy and mexiletine hydrochloride for Myotonic dystrophy type 1, showcased how these drugs ameliorate disease phenotypes. These findings enhance our understanding of each drugs’ mechanisms of action at the molecular level. Despite these promising results, the complexity of muscle dystrophies, involving multiple tissues, suggests a need for future models that incorporate a broader spectrum of relevant cell types. Enhancing the cellular complexity of these models could further refine their accuracy and utility in drug screening applications, promising more targeted and effective therapeutic interventions for these debilitating conditions. This study sets a robust foundation for the use of completely scaffold- and biomaterial-free biofabricated *in vitro* models in muscle dystrophy research, potentially other tissues and diseases, offering a promising platform for both mechanistic studies and the development of therapeutic strategies.

## Supporting information

Dystrophy Model - EvolvedBio - Supplementary Files

## Acknowledgement

We express our gratitude to SPARC Molecular Analysis Center, The Hospital for Sick Children, Toronto, Canada for conducting the proteomic analysis; Robarts Research Institute, Western University, London, Canada for performing the histology assessment. All other experiments were performed at Velocity Incubator, Waterloo, Canada. We appreciate the assistance from staff members at these institutions.

## Conflict of Interest

The authors declare no conflicts of interest. All research expenses were covered by Evolved.Bio.

## Data Availability Statement

The mass spectrometry proteomics data have been deposited to the ProteomeXchange Consortium via the PRIDE partner repository with the dataset identifier PXD052728. The custom Python scripts used in analyzing the data are available on GitHub at https://github.com/Evolved-Bio/invitroMuscleDystrophyAnchoredCellSheet. Any other data used in the study but not included here is available upon request.

## Declaration of use of generative AI

During the preparation of this work, the authors used Open AI’s ChatGPT and Anthropic’s Claude in order to grammatically edit the manuscript and debug the Python scripts. After using these tools, the authors reviewed and edited the content as needed and take full responsibility for the content accuracy.

## Notes

### Competing Interest Statement

The authors have declared no competing interest.

