## Supplementary material for "*in vivo*-like Scaffold-free 3D *in vitro* Models of Muscular Dystrophies: The Case for Anchored Cell Sheet Engineering in Personalized Medicine": Dystrophy Model - EvolvedBio - Supplementary Files: Dystrophy Model - EvolvedBio - Supplementary Information.pdf

**Supplementary File 1.** Proteomics expression values (normalized total precursor intensities) for the tested groups and their replicates.

**Supplementary File 2.** Significantly up- and down-regulated proteins when comparing different groups through differential expression analysis.

**Supplementary File 3.** Pathway enrichment analysis performed using g:Profiler for up- and down-regulated proteins when comparing different groups.

**Supplementary File 4.** List of proteins in each of the 6 clusters when comparing drug treated groups with their untreated counterparts.

**Supplementary File 5.** Statistical analysis contrasting the pairwise comparisons of different groups using Violin plots.

**Supplementary Video 1.** Differences in physical properties of the 3D anchored muscle fibers in DD and MD groups compared to HC.

**Supplementary Video 2.** Effect of treatment with mexiletine hydrochloride on physical properties of anchored 3D muscle fibers in MD group.

### Supplementary Figures

**Supplementary Figure 1.** Differentiation of skeletal muscle cells in 2D culture. Culture was started using $5 \times 10^5$ cells of each group in growth medium on day 1. On day 3, while all groups were near confluent, medium was switched to differentiation medium. Differentiation medium was refreshed every other day till day 8. Different cell types showed different morphologies before differentiation.

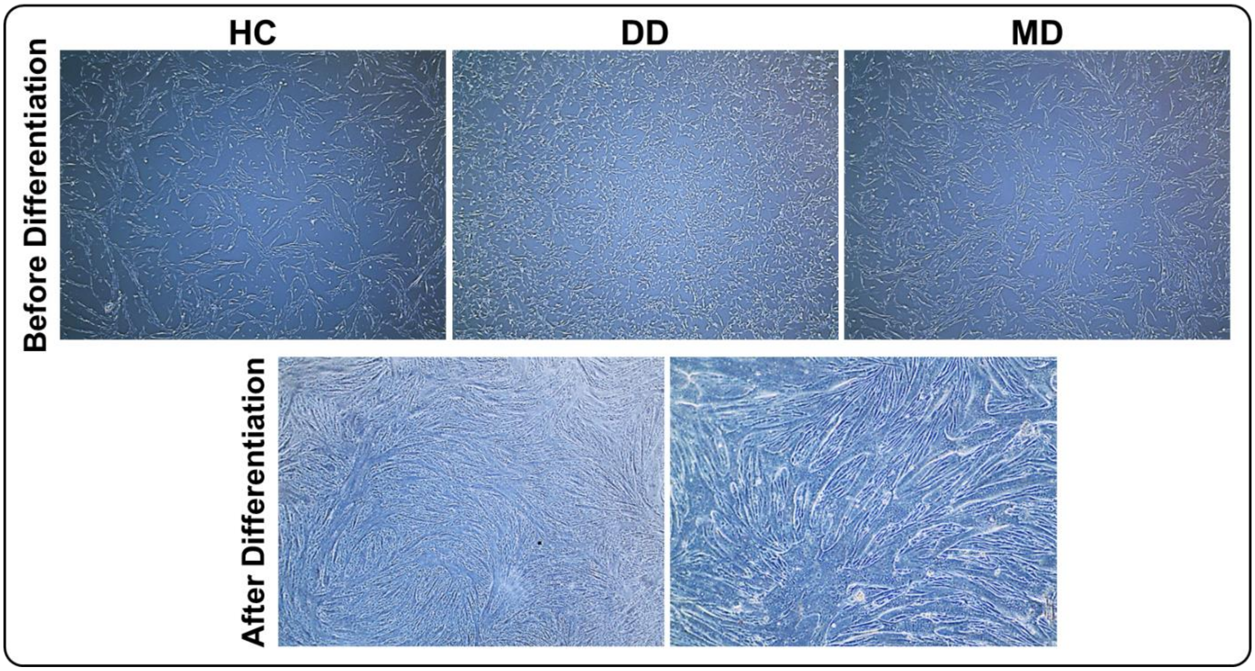

**Supplementary Figure 2.** Protein-protein interaction network made using STRING tool (https://string-db.org/) for proteins in cluster number 4 of **Figure 6a** (a total of 1066 proteins) of DD group treated with 1 mg/mL creatine. Majority of the proteins in this cluster belong to Extracellular Region (Red), Cell Junction (Purple), or Mitochondrion (Green) in cellular composition gene ontology analysis.

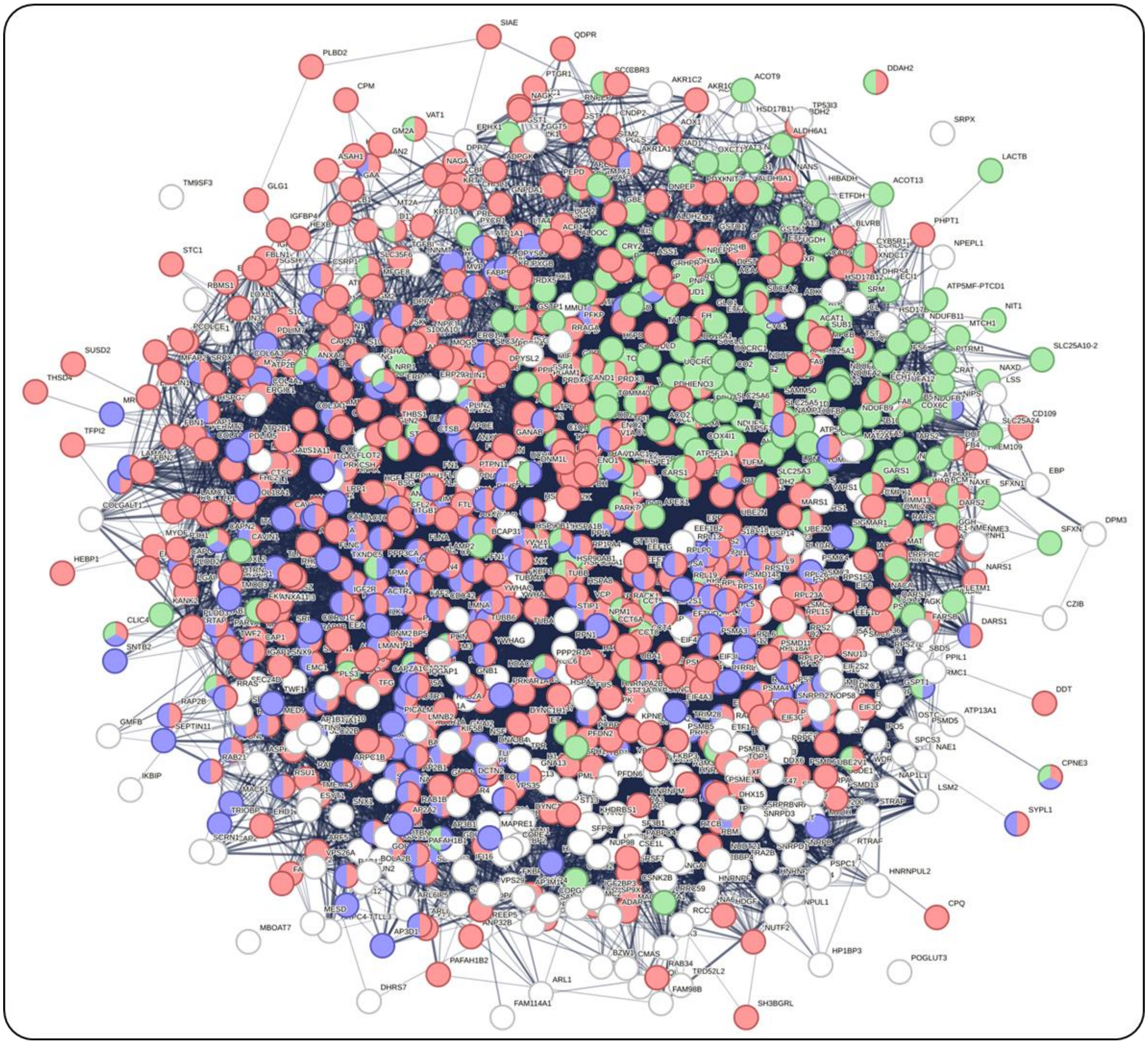

**Supplementary Figure 3.** Protein-protein interaction network made using STRING tool ([https://string-db.org/](https://string-db.org/)) for proteins in cluster number 4 of **Figure 6b** (a total of 1269 proteins) of MD group treated with 2 mg/mL mexiletine hydrochloride. Majority of the proteins in this cluster belong to Extracellular Region (Red), Cell Junction (Purple), or Mitochondrion (Green) in cellular composition gene ontology analysis.

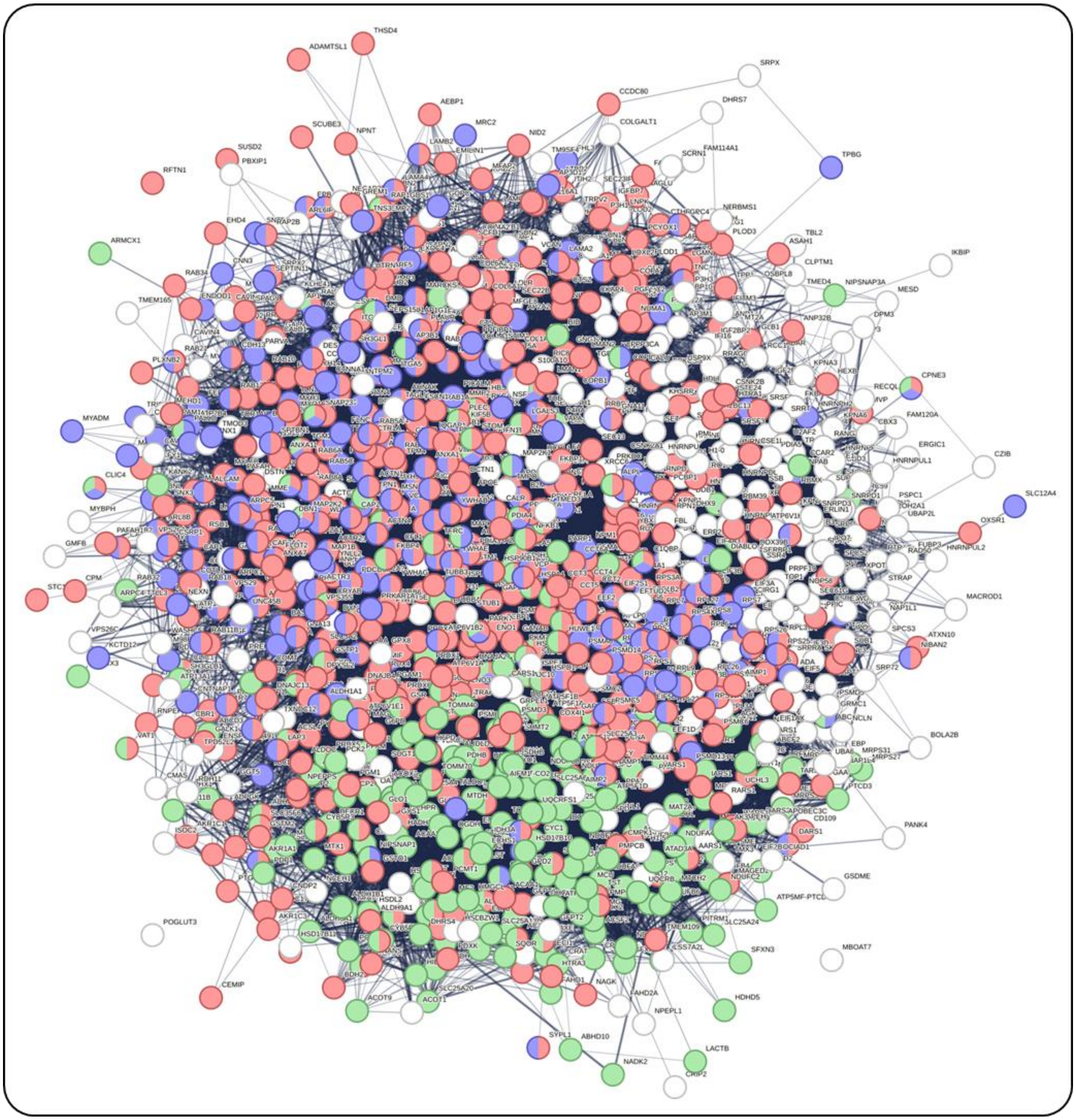

**Supplementary Figure 4.** Volcano and scatter plots showing the effect of treating **a)** DD and **b)** MD groups with higher drug concentrations. DD group was treated with 10 mg/mL creatin and MD with 20 mg/mL mexiletine hydrochloride. High similarities are observed between the two cases. Similar to lower concentration treatments, a high number of proteins were up- or down-regulated, when drug treated conditions were compared with their counterpart not treated with the drug. Comparison of the drug-treated DD and MD groups with the HC group also showed the formation of the same 6 distinct clusters, with the majority of the proteins located in clusters 3 and 4 along the diagonal. Some proteins exhibited minor changes in clusters 2 and 5, while others showed more substantial differences in clusters 1 and 6. L and H stand for low and high concentrations of drugs in each case, respectively.

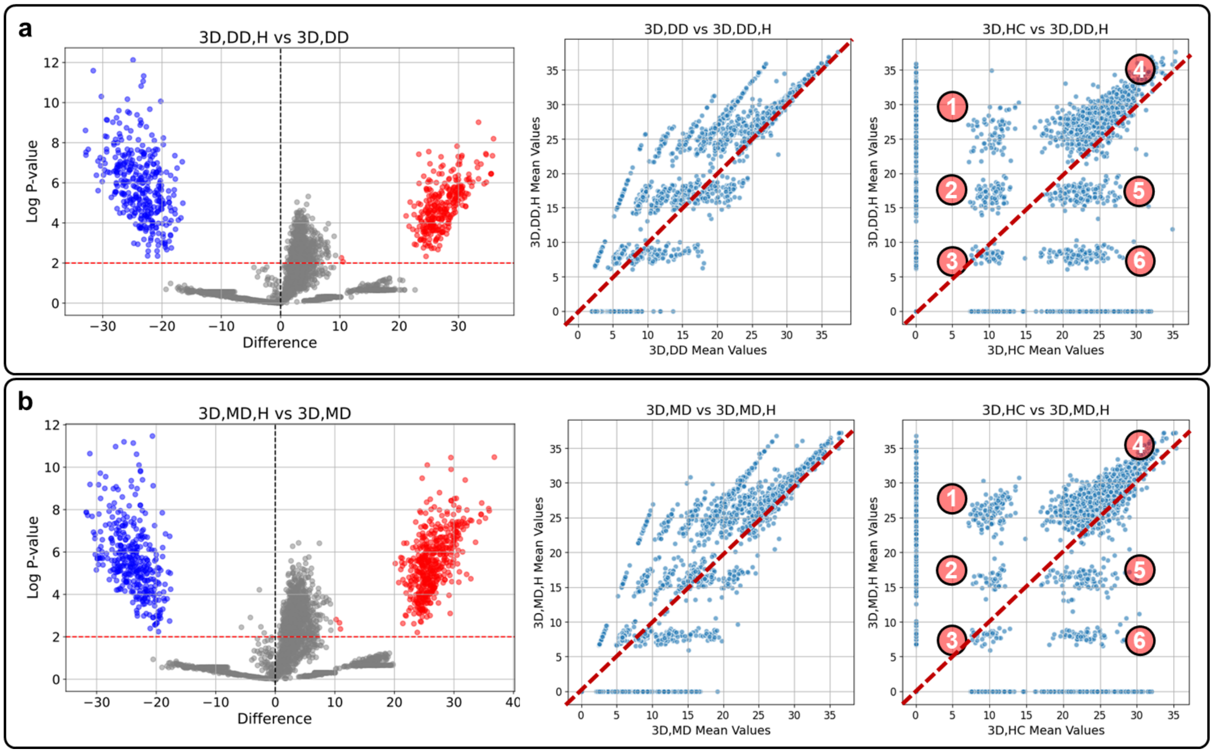

**Supplementary Figure 5.** Effect of drug concentration in each case using **a)** Scatter plots, **b)** Rank-Abundance plots and the Shannon Diversity Index showing diversity and richness of proteins, and **c)** Violin plots. In scatter plots had 9 separate clusters, three of which positioned on the diagonal, indicating high similarity between the two treatment concentrations while the other 6 showed varied levels of differences. The overall change in patterns in Rank-Abundance plots were similar, but Low concentration of creatine treatment in DD group showed earlier termination compared to the MD group where High concentration of mexiletine hydrochloride showed early termination. This is consistent with lower number of proteins observed under the diagonal of the scatter plots in DD group. In Violin plots, distribution of fold changes in protein expressions were shown when different groups were compared. Both low (L) and high (H) concentrations of drugs showed similar effects by making the plots wider and taller, and an upward shift.

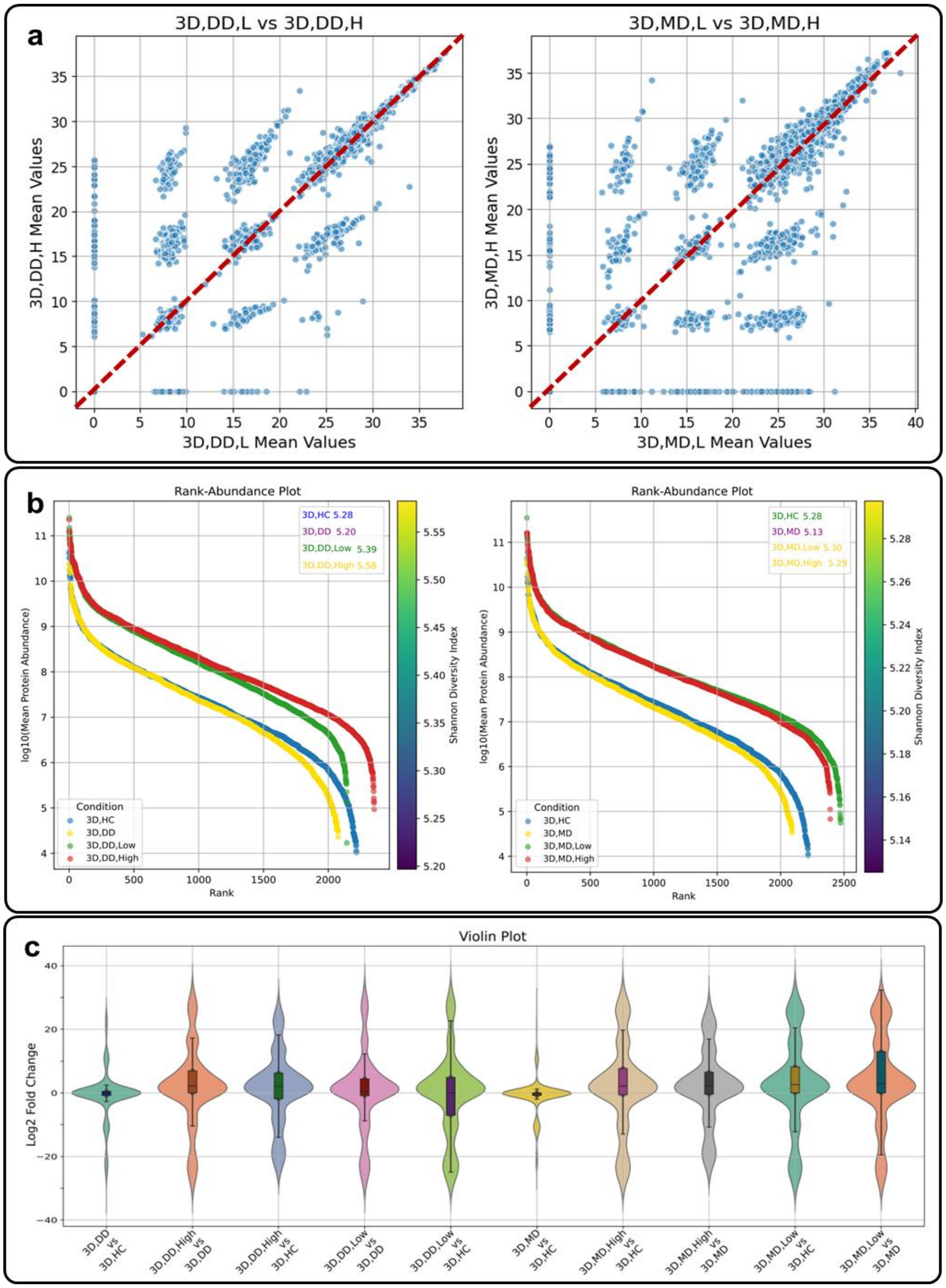
